# Trace_y: Software algorithms for structural analysis of individual helical filaments by three-dimensional contact point reconstruction atomic force microscopy

**DOI:** 10.1101/2023.07.05.547812

**Authors:** Wei-Feng Xue

## Abstract

Atomic force microscopy (AFM) is a powerful and increasingly accessible technology that has a wide range of imaging applications. AFM is capable of producing detailed three-dimensional topographical height images with a high signal-to-noise ratio. This is a key capability of AFM, which enables the structural features of individual molecules to be studied without the need for ensemble averaging, and could offer structural analysis applications where heterogeneity of molecular populations, structural variations between individual molecules, or population distribution properties in general, hold important information. Here, a software tool called Trace_y designed to reconstruct the three-dimensional (3D) surface envelopes of individual helical filament structures from topographical AFM height images is presented. Workflow using Trace_y is demonstrated on the structural analysis of individual helical amyloid protein fibrils where the assembly mechanism of heterogeneous, complex and diverse fibril populations from a single amino-acid sequence due to structural polymorphism is not fully understood. The software and the workflow presented here allows the structural information encoded in topographical AFM height images to be extracted and understood as 3D contact point clouds. This approach will facilitate the use of AFM in structural biology, in integrated methodologies together with other structural analysis tools to understand the structures and behaviours at individual molecule level.

## INTRODUCTION

Atomic force microscopy (AFM) is a powerful multimodal technique with imaging applications ranging from arrays of living cells to single molecules ^1^. AFM is an increasingly accessible method that has been used to study molecular dynamics ^2^ and molecular responses to force ^3^. It is also used for nanoscale imaging of biological molecules and structures in air or in liquid (e.g. ^4–6^). AFM topographical imaging operates by sensing the physical contact between the probe, typically consisting of a sharp conical tip attached to a cantilever, and the specimen surface, to generate a three-dimensional (3D) height map of the sample surface ^7^. While the imaging resolution of AFM is limited by the physical size of the AFM probe tip, molecular images acquired using AFM operating in non-contact mode with chemically functionalised probe tips is capable of reaching atomic resolution on samples of small planar molecules ^8^. Imaging resolution using un-modified probe tips can also be enhanced to sub-nanometres through recent image analysis advances ^9^. For structural biology applications, AFM offers several capabilities complementary to techniques such as cryogenic electron microscopy (cryo-EM) and X-ray crystallography. These include the ability of imaging in ambient conditions and in liquid environment, allowing imaging of biological samples such as live cells in a close to native environment ^10–12^. A key, sometimes overlooked, capability of AFM is that topographical images acquired by AFM can achieve a high signal-to-noise ratio, which enables structural features of individual molecules to be studied without the need for ensemble averaging ^13, 14^. This means that relevant structural information can be gained potentially from a single observation of a molecule on a single image. Therefore, this capability could offer new exciting opportunities in elucidating rare molecular structures in complex, heterogeneous samples and offer new information in the cases where the heterogeneity of molecular populations or the structural variations between individual molecules hold important biological relevance.

Amyloid fibrils represent a group of filamentous protein structures where the heterogeneity of the complex and mixed amyloid populations as well as the structural variations between the individual fibrils or even within individual fibrils hold relevant information regarding their biological impact ^15^. Amyloid fibrils are typically ordered helical nano-structures around 10 nm wide and between few nm to several microns long. They are assembled from monomeric proteins or peptide precursors. Some amyloid structures are associated with devastating neurodegenerative disorders including Alzheimer’s and Parkinson’s Diseases ^16–20^. In contrast, some amyloid fibrils have been identified to fulfil essential biological functions, providing scaffolding, adhesion and protection for organisms from bacteria to fungi, and more recently discovered to provide regulatory control of memory ^21^ and for viral infection ^22^. The formation of these functional amyloid fibrils is assumed to be tightly controlled. In contrast, the assembly of pathologically associated amyloid, even from a single type of precursor, can result in highly heterogeneous molecular mixtures, as well as a spectrum of different individual filament structures called polymorphs. Thus, it is clear that amyloid structural polymorphism is common for disease associated amyloid and this polymorphism phenomenon has been suggested to be a property differentiating disease associated amyloid from functional ones ^15^. AFM imaging has previously been used to characterise the morphological features of individual fibrils in a sample (e.g. ^13, 23–25^) and the size distributions of amyloid populations (e.g. ^26–30^). Recent advances in cryo-EM and solid-state nuclear magnetic resonance spectroscopy (ssNMR) have provided atomic-detailed structures showing multiple polymorphic forms for fibrils formed of identical precursors ^15^. However, to resolve the extent of the polymorphic diversity and the roles polymorphs play in biology, one must be able to determine the structures of individual amyloid fibril particles, and monitor their individual interactions in a complex and heterogeneous population to sufficient detail so that the diverse amyloid assembly landscapes, their polymorphic extents and biological properties linked to the heterogeneous populations can be mapped.

Here, an image analysis software Trace_y (https://github.com/wfxue/Trace_y), featuring a collection of algorithms for 3D contact point reconstruction (CPR) ^14^ and structural analysis of individual helical filaments, is presented. The individual filament structural analysis workflow using Trace_y is demonstrated here on the analysis of helical amyloid protein fibrils formed from the 42 residues long Amyloid β 1-42 peptide (Aβ_42_) associated with Alzheimer’s disease, where heterogeneous, complex and diverse fibril populations can assemble from a single monomer amino-acid sequence in the same sample due to structural polymorphism. Individual filament level structural analysis approach through 3D CPR-AFM, recently developed to utilise the high signal-to-noise topographical imaging capability of AFM ^13, 14, 31^, has the potential to tackle the challenges presented by the structural analysis of complex and heterogeneous samples such as polymorphous amyloid fibrils ^32^. The Trace_y software and the workflow for individual filament structural analysis presented here also demonstrate that the three-dimensional information encoded in AFM height images can be understood as 3D point clouds, which is a type of spatial information that could be integrated with other structural analysis methodologies ^33^.

## RESULTS

### Overview of the Trace_y workflow and its algorithms

The purpose of the workflow presented here is to recover the 3D structural information of individually observed helical filaments encoded in topographical AFM height images. **Figure 1** shows an overview of the workflow with a flowchart highlighting the key steps and the algorithms in the workflow with the example application to an individual fibril formed from the human Aβ_42_ sequence ^31^. The input for the workflow is an AFM height image where each pixel in the image contains the information of its x/y coordinate in the image lattice as well as the pixel value representing the recorded z-coordinate of AFM probe tip when intermittent contact between the tip and the molecular surface of the sample was detected. Due to the physics behind AFM, topographical height images have high vertical (z-axis) signal-to-noise ratio, with the noise typically in the order of sub-Ångströms range on current commercial instruments. One challenge of 3D structural reconstruction using AFM height image is that the recorded x, y and z-coordinates of the tip are not the actual tip-sample contact points ^14, 34^, which are not known. This issue is commonly referred to as the tip-sample convolution artefact. An important step in the workflow, therefore, is to deconvolute the image to recover the actual tip-sample contact points. Once the coordinates for tip-sample contact points are recovered, full 3D surface envelopes can be rendered for structures with helical symmetry, despite that only the top surface of the sample is accessible to the AFM tip, because structural information on the whole of a helical surface can be gained from a single direction of observation perpendicular to its helical axis due to their periodic twist around the helical axis. This requires the knowledge on the location of the helical axis, the handedness of the twist and the pitch of the helical structure, all of which can be gained or estimated from the height images (**Figure 1c-e**). The output of the workflow is a 3D contact point cloud of the surface of the helical filament, which can be further visualised as a 3D surface envelope model of the filament structure. The algorithms performing the key steps of the Trace_y workflow are explained in the sections below and exemplified on a fibril formed from recombinant human Aβ_42_ (**Figure 1b-c**).

**Figure 1.**
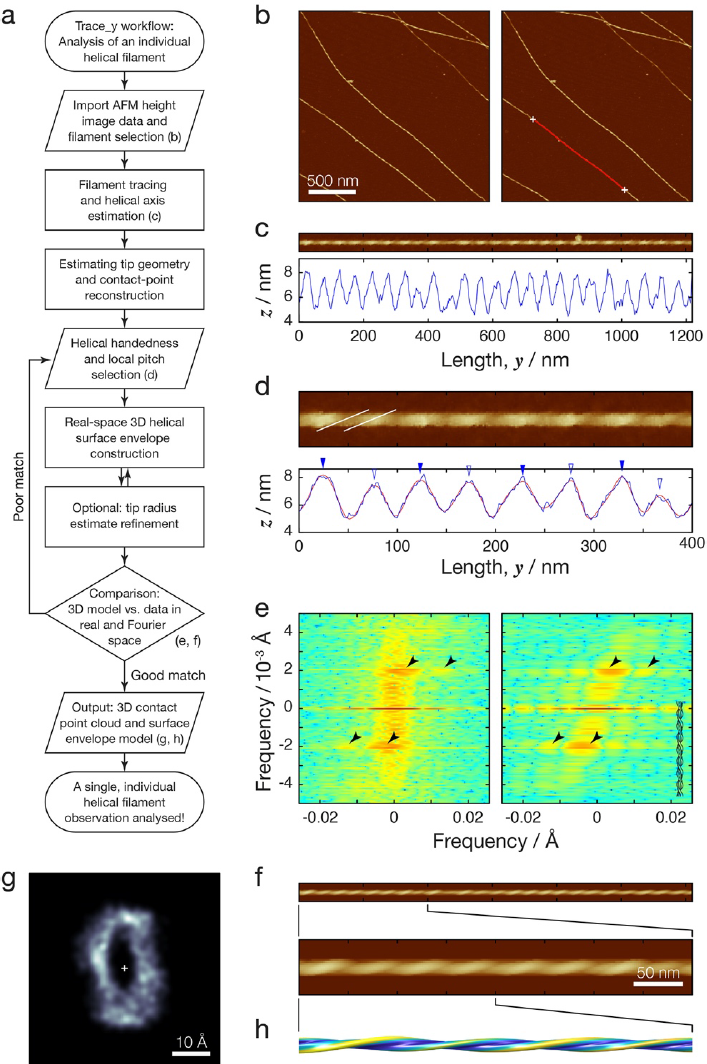
The Trace_y workflow demonstrated on an individual Aβ_42_ amyloid fibril. (a) Flowchart overview of the Trace_y workflow for analysing the structure of an individual helical filament. (b) Topographical AFM height images of helical twisted Aβ_42_ fibrils. The right image is identical to the left image, except with an example fibril and its central contour line highlighted (red line). The white crosses indicate user defined start and end point of the traced fibril segment. The scale bar represent 500 nm in both images. (c) Digitally straightened fibril image showing the fibril highlighted in (b) and its central line height profile. (d) A zoomed image showing the first 400 nm segment the same fibril shown in (c). Its central line height profile (blue line) is shown together with a digitally smoothed line (red line) with which the peak positions were determined (blue triangles). The filled blue triangles indicate reference locations with one helical pitch distance in between each filled blue triangles. (e) Power spectrum of 2D Fourier transform of the fibril image data is shown to the left compared with that calculated from a simulated topograph image shown to the right. The black arrows indicate identical locations in both power spectra and show major peak locations. (f) Topographic AFM height image simulated from the 3D fibril envelope model (h) of the same fibril in (b) together with a zoomed in image showing the first 400 nm segment. (g) The cross-sectional contact-point density map of the same highlighted fibril from (b) Scale bar indicate distance of 10Å.

### Individual filament tracing and estimation of the helical axis

A semi-automatic fibril tracing algorithm was developed to allow the user to select and to trace segments of individual filaments observed on AFM height images. Since the central contour line of a filament appearing in a height image typically corresponds to the projection of the filament’s helical axis on the x/y-plane, an accurate estimate of the central contour line of the selected filament is needed. Therefore, a curved Gaussian wall ^35^ model (Equation 1, Methods) was used and fit by least-squares method to regions of the height image (image squares with edges twice the length of filament’s apparent width) cropped along the coarse user-selected contours of the filament (**Figure 2**). This approach allows the x/y-coordinates of points on the central contour line of the filament to be determined with sub-pixel accuracy because the coordinates are estimated using information from the entirety of the local features rather than few individual pixels. The z-coordinate of the filament helical axis was subsequently estimated as half of the average peak height (for example, the peaks values at locations marked with triangles in **Figure 1d**). This simple estimate assumed a rigid fibril that rests on the support surface without substantial deformation during deposition and by the peak probing force during imaging.

**Figure 2.**
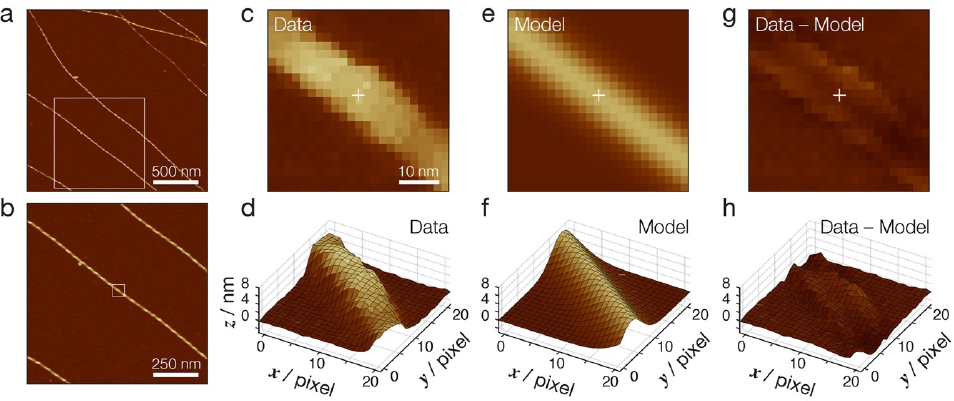
Tracing of individual filaments and estimation of helical axis coordinates by regression analysis using a curved Gaussian wall model. (a) Topographical AFM height images of helical twisted Aβ_42_ fibrils. The same 2×2 μm image as Figure 1b is shown. The white box indicate 1×1 μm magnified area shown in (b) and scale bar indicating 500 nm. (b) 1×1 μm magnified area indicated by the white box in (a). The white box in (b) indicate 22×22 pixel area shown in (c)-(f), and scale bar indicating 250 nm. (c) and (d) 22×22 pixel area indicated in the image data in (b) shown as a 2D height image and as a 3D surface, respectively. (e) and (f) Best fit curved Gaussian wall model of the same area indicated in the image data in (b) shown as a 2D height image and as a 3D surface, respectively. (g) and (h) Pixel level residuals between the image data and the best fit model shown as a 2D height image and as a 3D surface, respectively. In (c)-(h), white cross indicate an estimated point of helical axis x/y coordinates. Scale bar indicate 10 nm in (c), (e) and (g).

## 3D contact point reconstruction by image deconvolution

The 3D coordinates of tip-sample contact points were next reconstructed by modelling rigid-body interactions between the AFM probe tip and the sample filament using the image deconvolution strategy as previously described ^14^. First, the tip geometry was estimated. A rounded cone tip model made by the rotation of the half cross-section described by Equation (2) around the y-axis is used and the tip radius was estimated using a iterative approach where image dilation caused by the contact between the tip model and the circular cross-section of a featureless cylinder is matched to the experimentally observed dilation for the analysed filament. This procedure is illustrated by **Figure 3a**, where least-squares regression analysis is performed with a calculated convoluted cross-section (red lines and crosses) of a featureless cylinder (black) fit to the observed convoluted cross-section of the filament averaged through the length of the filament along its estimated helical axis (blue circles). This approach is able to provide a useful initial tip radius estimate using the helical axis coordinates as input since the cross-section of a helical structure with its cross-section rotating around its helical axis approaches a circular shape when averaged over the length of its helical axis. The initial tip radius estimate can be further optimised in a later optional step by minimising the difference between the image data of the filament and a simulated image calculated using the surface envelope model of the same filament. An important advantage of this sample feature-based blind tip radius estimation approach is that additional dedicated tip characterisation step, which can additionally blunt the tip during characterisation, can be avoided.

**Figure 3.**
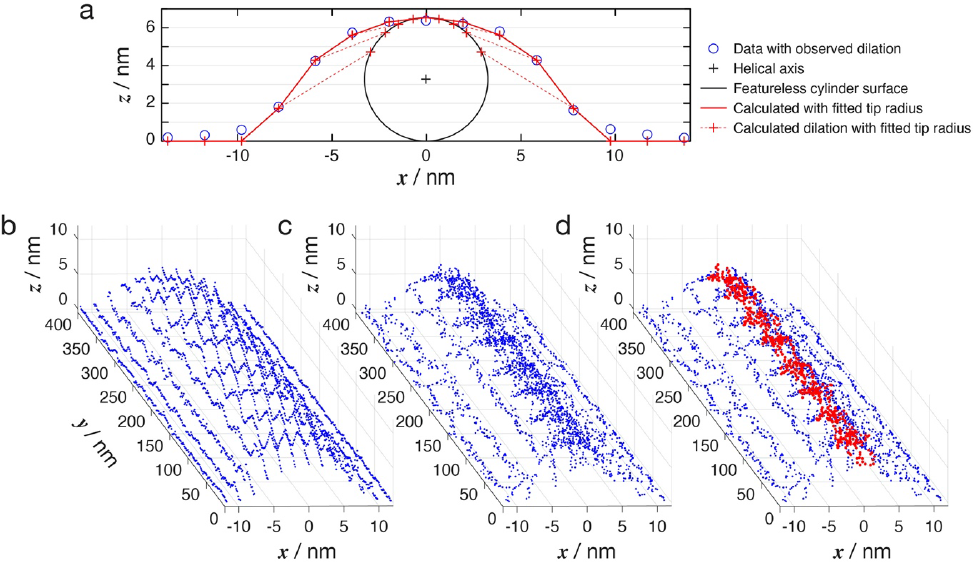
Cantilever tip radius estimation and contact point reconstruction result in a 3D point cloud that represents the tip-filament surface contact points. (a) Initial estimation of the tip radius by least-squares fit of calculated image dilation (red crosses, red solid line and red dotted lines) from a featureless cylinder (black line representing the cross-section of the cylinder) to observed average dilation along the fibril (blue circles). (b) The fibril image in Figure 1d shown as points in 3D space representing the pixels’ x/y/z coordinates. (c) The point cloud output of CPR showing the x/y/z coordinates of the points in (b) moved to estimated tip-filament surface contact points. (d) Contact points on the top of the helical filament subsequently used for construction of 3D surface envelop is show as red points.

Once the tip radius is estimated, deconvolution of the image data by CPR ^14^ was subsequently carried out to recover the 3D coordinates of tip-sample contact points (**Figure 3b-d**). This step outputs a point cloud containing the estimated tip-sample surface contact points by moving the 3D x/y/z-coordinates of the recorded tip positions off the x/y pixel lattice of the image to the appropriate sub-pixel locations estimated by CPR step. **Figure 3b** shows the AFM height image of a segment of the example Aβ_42_ filament where each pixel is shown as a point in 3D space. All of the points have their x/y coordinates in the pixel lattice of the image, and the pixel lattice along the x-axis is particularly visible from the viewing angle used in **Figure 3b**. **Figure 3c** demonstrates the tip-sample contact point cloud after image deconvolution by CPR where the data points are no longer bound to the x/y pixel lattice and instead echo the actual sample surface coordinates. The points that are localised at the top of the filament (red points in **Figure 3d**) were then selected to be used in subsequent steps to create a surface envelope model of the individual filament. In summary, the CPR step reconstructed the 3D contact points between the AFM tip and the filament, creating a point cloud that represents the top of the individual filament being analysed.

### Visualisation of the filament cross-section and construction of 3D surface envelope model

The point cloud data that represents the structure of the filament top obtained from the CPR step was next used to create a 3D surface envelope model of the filament. The helical properties of the individual filament being analysed were first determined by manual inspection, this is shown for the example Aβ_42_ filament in **Figure 1c-e**. Since topological AFM height images are not transmission images, only one side of the helical filament is observed. Hence, the helical handedness of the filament twist is typically readily visible by the tilt direction of the striation pattern (exemplified by the white lines in **Figure 1d**). For left-hand twisted filaments, the tilt of the pattern goes from bottom left to top right whereas the tilt of the pattern goes from bottom right to top left for right-hand twisted fibrils when the filaments image is aligned horizontally. The handedness can also be deduced by 2D Fourier transform of the filament image (**Figure 1e**). For vertically aligned filaments images, the main peaks in the 2D Fourier transform power spectrum image follow the same bottom left to top right or bottom right to top left tilt patterns relative to the image centre for left-handed or right-handed twists, respectively (arrow heads in **Figure 1e** for the example Aβ_42_ filament). The helical pitch of the filament can also be deduced from the z-height peak patterns of the central contour line of the filament image, and the peaks in the 2D Fourier transform of the filament image. In this case, a direct determination of the helical pitch by spotting repeating patterns in the z-height profile or estimating the magnitude of the tilt of the pattern in 2D Fourier transformed image may not be straight forward as the differences expected for different cross-sectional symmetries may be small. Therefore, a best guess of the cross-sectional symmetry and resulting helical pitch was used for the initial 3D model building, and the validity of this initial guess was checked by comparing simulated image and its 2D Fourier transform power spectrum with the image data of the filament in a subsequent step. This is illustrated for the example Aβ_42_ filament in **Figure 1d** where the helical pitch was estimated to be the peak-peak distances of every-other peak in the central line z-height profile of the filament (e.g. filled blue triangles in **Figure 1d**), possibly due to a C_2_ cross-sectional symmetry or a 2_1_-screw axis symmetry. This guess for the example filament was validated because the simulated image based on the 3D model constructed using the guessed pitch values (**Figure 1f**) matched well with the image data, and the location of the main peaks in the 2D Fourier transformed image matched with those in the 2D Fourier transformed power spectrum image of the data (arrow heads in **Figure 1e**).

Once the helical pitch and twist handedness is determined for the individual filament being analysed, the point cloud data representing the structure of the filament top is moved from its original AFM height image coordinate system to a helical coordinate system (**Figure 4a**) where the helical axis of the filament is aligned with the y-axis and the x/z-axes (the cross-sectional plane) rotates with the same rate along the helical axis as the helical filament being analysed. In this case, the x/z-plane rotates with a θ rotation angle that is locally estimated, referenced onto the same peaks used to calculate the local helical pitch values (filled blue triangles in **Figure 1d**). For the example Aβ_42_ filament, the θ rotation angle at any given y-coordinate along the helical axis is shown in **Figure 4b** where the blue crosses are the local reference points at full integral number of complete turns around the helical axis, and the red line represents the interpolated θ values along the helical axis by cubic splines interpolation. The cross-sectional envelope of the helical filament was then visualised on the helical x/z plane when the point cloud was registered onto the rotating x/z-frame. **Figure 1g** shows the visualisation of the cross-sectional envelope of the example Aβ_42_ filament as a density map of the contact points in the x/z-plane constructed using a non-parametric kernel density estimator ^36^. The spatial resolution of this density map was subsequently estimated by a Monte Carlo (MC) approach where the points in the point cloud were randomly split into two equal sized sets. The density map of each set of points were separately constructed (an example pair is shown in Figure 4c) and their Fourier ring correlation were evaluated using the ½-bit information criterion ^37^. For the example Aβ_42_ filament shown in **Figure 1g**, evaluated using 500 MC cycles, the resulting spatial resolution estimate was around 13.1Å (**Figure 4d**). Finally, the 3D surface envelope of the whole of the filament was constructed by fitting an average cross-sectional envelope constructed by a cubic spline curve in the x/z-plane to the point cloud data. A moving window approach to select only points local to each evaluated y-coordinate (within ±1 helical pitch) were used to construct cross-sectional envelopes that takes into account local structural variations as previously described ^14^. The final 3D surface envelope of a 400 nm segment of the example Aβ_42_ filament is shown in **Figure 4e** and **Figure 4f**. The filament surface was coloured by its perpendicular distance to the helical axis in **Figure 4e** to highlight its helical properties and by its local twist angle per unit length in **Figure 4f** to highlight the structural variations in its twist along this single individual filament.

**Figure 4.**
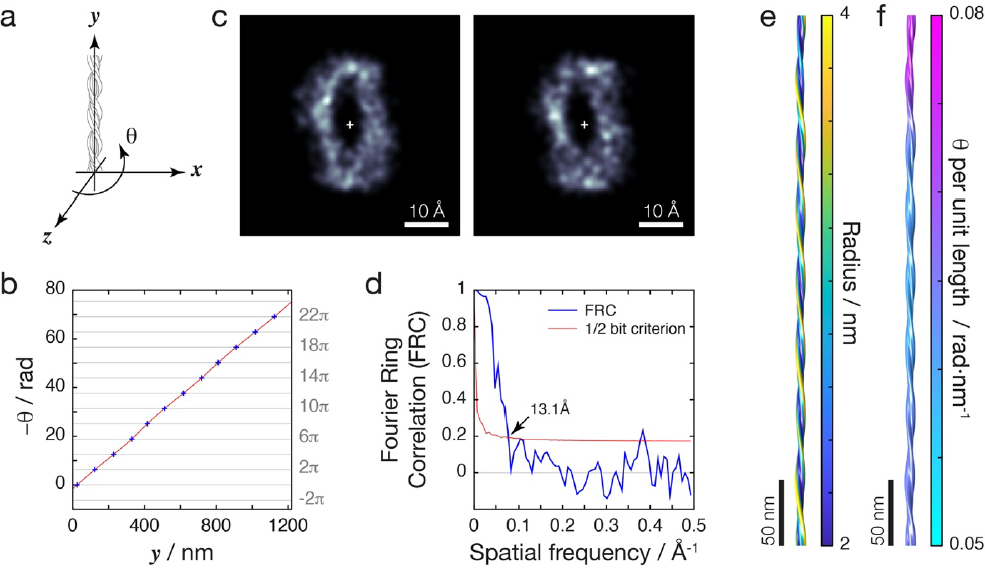
Construction of filament 3D surface envelope by point cloud registration in a helical coordinate system. (a) Schematic of the helical coordinate system used to align contact points in a point cloud representing a helical filament. The x/z plane of this coordinate system is set to rotate around the helical axis aligned to the y-axis with the same twist angles 8 as observed for the filament. (b) The local twist angle 8 (blue cross) of the example filament shown in Figure 1c measured using its height profile (e.g. filled blue triangles in Figure 1d). Red line represents the 8 function along the whole filament constructed by spline interpolation with the measurement points (blue cross). (c) The cross-sectional contact-point density map of the same example filament. An example density map pair generated by randomly selected half-sets of the contact point cloud used for FRC measurement is shown. (d) The spatial resolution of the contact-point density map shown in Figure 1g estimated by FRC (blue line) and the ½ bit information criterion ^37^ (red line with the FRC crossing point indicated by the arrow). (e) 3D surface envelope model of the example filament (the first 400 nm segment as shown in Figure 1d) with surface colour indicating distance to helical axis. (f) The same 3D surface envelope model of the example filament as (e) but with colour indicating helical twist mapped to its surface to highlight local helical structural features.

## DISCUSSION

Atomic force microscopy has recently embarked on a renaissance in terms of its application in imaging of biological/soft materials and in molecular structural biology, driven by the recent developments in high-speed AFM instrumentation ^38^ and several new emerging image analysis algorithms ^9, 14, 39–42^. Despite the limitations of the AFM physics due to the nanometre size of the probe tip and that only the surface features of molecules of interest are probed, AFM is capable of time-resolved imaging with ∼100 ms temporal resolution in aqueous conditions, allowing the dynamics of molecules to be studied under ambient conditions. AFM is also capable of high signal-to-noise topology imaging with typically sub Ånström noise and ∼1Å resolution in the z height-axis. This feature allows for structural analysis of individual molecules without cross-molecule averaging ^13, 14, 31^. Therefore, this allows the structural variations between individual molecules in a population or within the same molecular assembly to be resolved. Together, these two notable features of AFM fill an important gap in the type of molecular structural information that can be gained, and creat an unique niche for AFM in structural biology where methods such as cryo-electron microscopy, nuclear magnetic resonance spectroscopy and X-ray crystallography all provide ensemble averaged structural information. Here, the high signal-to-noise topology imaging feature of AFM is exploited to allow 3D structural studies of individual helical filaments.

The Trace_y software and the workflow based on 3D CPR-AFM presented here is a post-experiment analysis approach. As such, it is widely applicable for the analysis of topographical AFM height images. Importantly, the application of Trace_y on helical amyloid fibrils demonstrated here show unequivocally that 3D structural reconstruction is possible using AFM data as input. The CPR approach allows this by unlocking the 3D information encoded in the images and show that this information can be understood and visualised as 3D contact point clouds. Contact point clouds can be quantitatively analysed once aligned to an appropriate and meaningful coordinate system and/or to each other, this is a process also referred to as point cloud registration. Indeed, point cloud registration is a principal problem that features prominently in pattern recognition and computer vision fields. In the case of Trace_y, the current presented version of the 3D reconstruction workflow and the algorithms that underpin the key steps of the workflow establishes many opportunities for further developments, refinements and improvements at each and every step by the bio-AFM community. For example, refinements to the helical axis estimate and subsequently the alignment of the cross-sectional axes will improve the overall quality of the 3D point cloud in representing the surface envelope of the helical structures of interest. Integrative approaches involving spatial 3D data sources from cryo-EM, molecular PDB models from NMR or X-ray crystallography studies, or predicted molecular models (e.g. by AlphaFold ^43^) can further extend the use of the 3D point cloud data and provide models by which the point clouds can be compared to or registered onto. In addition, it may be possible to map chemical or nanomechanical information onto the 3D surface envelopes (e.g. using analogous approach as demonstrated in **Figure 4e, f**) reconstructed from multi-modal AFM-IR, AFM-Raman or nanomechanics AFM experiments. Therefore, these opportunities may make it possible to distinguish between different conformational or ligand binding states of biological molecules or complexes, facilitating the analysis of protein structure and function to high details on a single, individual molecule level.

## MATERIALS AND METHODS

### AFM specimen preparation

The 3D contact-point reconstruction AFM (CPR-AFM) workflow to analyse the 3D surface envelopes of individual helical filaments using Trace_y is demonstrated on amyloid fibrils formed from the 42 residues long Amyloid β 1-42 peptide (Aβ_42_). Helical filaments formed from Aβ_42_ is associated with neurodegenerative diseases such as Alzheimer’s disease (AD), in which extracellular deposits known as plaques containing Aβ_42_ amyloid fibrils are found in brains of patients ^44^. Here, fibrils were formed *in vitro* from recombinant Aβ_42_ ^45^ in 20mM HEPES buffer under quiescent conditions at pH 7.4 and 37°C ^31^. The fibrils were deposited onto freshly cleaved mica discs as previously described ^31^.

### Topographical AFM image data acquisition

Aβ_42_ amyloid fibrils were imaged by AFM using a Multimode 8 SPM with a Nanoscope V controller (Bruker) operating under peak-force tapping mode as detailed previously ^13^. Briefly, ScanAsyst silicon nitride probes (Bruker) with nominal tip radius of 2 nm and nominal spring constant of 0.4 N/m were used. Topographical height images with a scan size of 4 x 4 µm and 2048 x 2048 pixels or 8 x 8 µm and 4096 x 4096 pixels were collected maintaining a pixel density of 0.512 pixels/nm. A scan rate of 0.305 Hz was used for the 4 x 4 µm and 0.2 Hz for the 8 x 8 µm scans. The fibrils were probed gently using ∼400 pN of peak-force, with a noise threshold of 0.5 nm and the Z limit was of 1.5 µm were used.

### Initial image processing

Nanoscope analysis software supplied with the Multimode 8 SPM instrument (updated to version 1.5, Bruker) were used for the initial processing of the z-height image data. Line flattening were performed to remove tilt and scanner bow by fitting a 3rd-order polynomial baseline to each scan line of the z-height channel image. For background flattening, the thresholding option in Nanoscope analysis was used and the thresholding level was manually adjusted prior to the polynomial fit. The height channel of the data was then imported into Trace_y for the subsequent steps of the workflow.

### Algorithm implementation and image analysis

The algorithms in the Trace_y workflow were implemented in Matlab (2022a, The MathWorks, Natick, MA) and include graphical user-interface for various steps in the workflow that requires user inputs. Equation (1) describes the Gaussian wall model ^35^ used to determine a point’s coordinates *x̂* and *ŷ* along the central filament contour.

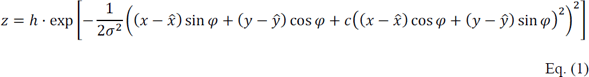

In Equation (1), *h* is the height of the Gaussian wall, σ is the standard deviation of the Gaussian cross-section, φ is the angle of the filament on the x/y-plane and *c* is the curvature parameter. The 3D reconstruction of tip-sample contact points were performed by image deconvolution as previously described ^14^. For the deconvolution procedure, a rounded cone tip model made by rotation of the half cross-section described by Equation (2) around the y-axis is used.

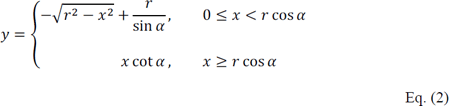

In Equation (2), *α* is the half-angle of the tip cone and was set to 18° here (estimated from the tip geometry information provided by the manufacturer). The *r* parameter is the tip-radius. Software package with stand-alone executable of Trace_y was prepared using Matlab Compiler (2022a) and can be downloaded from the GitHub (https://github.com/wfxue/Trace_y).

## ACKNOWLEDGMENTS

Help with the software development and insightful discussions by Liisa Lutter, and acquisition of the Aβ_42_ amyloid image data used here as examples by Liam Aubrey are gratefully acknowledged. Testing and helpful comments provided by the members of the Xue and the Louise Serpell research groups are also gratefully acknowledged. The AFM image examples used is acquired on an instrument supported by funding from the Biotechnology and Biological Sciences Research Council (BBSRC), UK grants BB/J008001/1, BB/M02427X/1 and BB/S003312/1

